# Quantification of inter-sample differences in T cell receptor sequences

**DOI:** 10.1101/128025

**Authors:** Ryo Yokota, Yuki Kaminaga, Tetsuya J. Kobayashi

**Affiliations:** Institute of Industrial Science, the University of Tokyo, Tokyo, Japan; Department of Electrical Engineering and Information Systems, Graduate School of Engineering, the University of Tokyo, Tokyo, Japan; PRESTO, Japan Science and Technology Agency (JST), Saitama, Japan

**Keywords:** T cell, TCR repertoire, inter-sample comparison, pairwise sequence alignment, sequence dissimilarity, manifold learning, Jensen-Shannon Divergence

## Abstract

Inter-sample comparisons of the T cell receptor (TCR) repertoire are crucial for gaining a better understanding into the immunological states determined by different collections of T cells from different donor sites, cell types, and genetic and pathological backgrounds. As a theoretical approach for the quantitative comparison, previous studies utilized the Poisson abundance models and the conventional methods in ecology, which focus on the abundance distribution of observed TCR sequences. However, these methods ignore the details of the measured sequences and are consequently unable to identify sub-repertoires that might have the contributions to the observed inter-sample differences. In this paper, we propose a new comparative approach based on TCR sequence information, which can estimate the low-dimensional structure by projecting the pairwise sequence dissimilarities in high-dimensional sequence space. The inter-sample differences are then quantified according to information-theoretic measures among the distributions of data estimated in the embedded space. Using an actual dataset of TCR sequences in transgenic mice that have strong restrictions on somatic recombination, we demonstrate that our proposed method can accurately identify the inter-sample hierarchical structure, which is consistent with that estimated by previous methods based on abundance or count information. Moreover, we identified the key sequences that contribute to the pairwise sample differences. Such identification of the sequences contributing to variation in immune cell repertoires may provide substantial insight for the development of new immunotherapies and vaccines.

## 1 INTRODUCTION

The development of high-throughput sequencing with next-generation sequencers has provided new opportunities to quantify T cell receptor (TCR) repertoires and to compare their differences among different cell types, organisms, and pathological samples. Such information is indispensable for quantitatively understanding the immunological state of organisms that is shaped by the collection of immune cells. Moreover, the detailed information of TCR repertoires, especially that of inter-sample differences, is anticipated to significantly promote the development of immunotherapies and vaccines (Hou et al., 2016). To this end, several theoretical methods have been proposed to quantify sample differences by focusing on the count (abundance) distribution of unique TCR sequences in a repertoire (Greiff et al., 2015; Laydon et al., 2015; Hou et al., 2016). Poisson abundance (PA) models are among the recently developed methods based on the hierarchical Bayesian inference algorithm, which estimates the parameters of the models from experimental data and defines the inter-sample difference according to the deviation of the estimated parameters. This method overcomes the substantial sampling fluctuations derived from the huge diversity in TCR repertoires, and provides a stable result related to the inter-sample distances on the basis of statistical interpretations. For example, Rempala et al. (2011) used a bivariate Poisson log-normal (BPLN) model to classify eight different samples of the following sample conditions: donor sites, types of T cells, and the genetic backgrounds of different mouse lines. Guindani et al. (2014) used a Poisson Dirichlet process to classify the types of T cells (i.e., conventional and regulatory T cells). Besides the above examples, other variations of PA models have been proposed for sample classification based on the measurement of TCR diversity (Sepúlveda et al., 2010; Greene et al., 2013).

Although these PA models can successfully quantify the inter-sample distances, they are also associated with a major drawback in that some of the sequence information for each sample is lost since these models focus only on the count distribution. This loss of information has hampered the ability to determine the characteristic sequences of each sample, which is a requisite for further investigations of the source of the difference by, for example, evaluations of the interaction with microbial peptides (Aas-Hanssen et al., 2015) and the simulation of TCR crystal structures (Klausen et al., 2015).

As an alternative method, we can extract and count the overlapping sequences between two samples or among many samples. However, the possible sequence space of the TCR repertoire is at least as large as 10^15^ (Davis and Bjorkman, 1988), and therefore the measured sequences can only sparsely cover the entire space. This sparsity substantially reduces the chance to observe the same sequence in two samples. Thus, by focusing on the overlaps, it is only possible to detect the public sequences that appear very frequently among the samples. Moreover, even if no overlapping among the sequences is detected, it is not possible to judge whether this occurs because the two repertoires cover quite different subspaces of the sequence space or because the repertoires cover the same subspace but show no overlapping by chance simply owing to the sparsity of the coverage. This difference can be determined by exploiting the information of inter-sequence differences in the repertoires.

To address this problem, we developed a new method for the quantitative comparison of TCR repertoires, by focusing on the sequence information in all samples, and estimating the low-dimensional structure (manifold) by projecting the high-dimensional inter-sequence relations, calculated from pairwise sequence alignments, onto a low-dimensional space. The methods for manifold estimation have been successfully applied in previous studies of virus evolution (Ito et al., 2011) and relationships of 16S rRNA gene sequences in bacterial genomes (Hughes et al., 2012) to extract the evolutionary pathways and interconnections of bacteria. Although manifold estimation has also been employed for evaluating the TCR repertoire, this was mainly used only for visualization purposes (Duez et al., 2016). However, the low-dimensional embedding of the original repertoire contains the information how the repertoires from different samples cover the possible sequence space. Therefore, by employing such information, it may be possible to detect a subset of sequences in the repertoire that has a major contribution to the inter-sample difference.

To quantitatively compare the embedded sequences, we estimated a probability density function of the sequence distribution in the low-dimensional space. This density estimation compensates for the sampling bias due to unseen sequences from the sparsity of the measured sequences. Finally, we quantified the inter-sample differences between the estimated density functions of the individual samples using the Jensen-Shannon divergence (JSD). This information-theoretic measure characterizes the difference between two distributions by the probability of observing either one by chance with random sampling from the others. Thus, this measure can effectively and quantitatively capture information on the existence of non-overlapping sequences between two repertoires. By extracting the sequences that show a major contribution to the information theoretic measures, the sequences most responsible for the inter-sample differences can be determined, which cannot be identified with previous approaches.

The paper is structured as follows. We first describe the experimental data adopted to test our method and the step-by-step data analysis procedure, including (i) quantification of sequence dissimilarity with the pairwise sequence alignment algorithm, (ii) evaluations of four different manifold learning methods for projecting the sequence distribution in low-dimensional space, (iii) adoption of the kernel density estimation algorithm (KDE) to quantify the sequence distribution, and (iv) quantification of the inter-sample differences and identification of the contributing sequences according to the JSD values of the distributions. To validate the applicability of our method, we apply a true dataset of TCR repertoires and demonstrate that similar inter-sampling clustering can be obtained by both our method and previous methods despite their use of different modalities (sequence and count, respectively) of a repertoire. We further evaluate the statistical significance of our results using a bootstrap algorithm to confirm the derived sample difference. Overall, we aim to demonstrate the advantages of our method to previous methods by capturing more complete and quantitative information on TCR repertoires. This method is expected to be of value for understanding variation of the immunological states to facilitate development of immunotherapies and vaccines.

## 2 MATERIAL & METHODS

### 2.1 Sequence data

In this study, we used a public dataset of TCR repertoires in mice published in Rempala et al. (2011). This dataset includes information of eight different TCR populations, which are classified according to donor sites, types of CD4+ T-cells, and the genetic backgrounds of mice. The CD4+ T-cells were collected and isolated either from the thymus or peripheral lymph nodes, which are labeled as ”1” and ”2”, respectively. In addition, these cells were categorized into either naive T-cells (TN) or regulatory T-cells (TR) in accordance with the presence or absence of Foxp3 expression. The two genetic backgrounds of mice are labeled as “wild type” and “Ep” in the original paper. Both groups showed strong restriction on rearrangement of V(D)J genes (i.e., the two *α*-chain rearrangements between J*α*2.6 and J*α*2 with a fixed V*α*2.9 segment and fixed *β*-chain V*β*14D*β*2J*β*2.6). The main difference between these groups is that the Ep mice were backcrossed with mice that express transgenic class 2 major histocompatibility complex molecules bound to a single “Ep” peptide (Pacholczyk et al., 2006). Thus, Ep mice are expected to show a more restricted TCR repertoire than wild-type mice. To evaluate the diversity of TCR repertoire, the complementarity determining region 3 (CDR3) of TCR*α* chains were sequenced and amplified. Further details on this dataset are described in Section 4 of Rempala et al. (2011).

### 2.2 Data analysis procedure

To date, no sufficient and effective method for comparing TCR repertoires among different samples has been established, which is mainly due to the enormous complexity and diversity of TCR sequences (Hou et al., 2016). In ecology, the diversity in a population is conventionally measured with metrics of species abundance’ between pairs of samples. However, these methods generally rely on observational data of species abundance counts, and can therefore be vulnerable to sampling bias. One widely used typical measure to quantify biological diversity is through a dissimilarity metric such as the Bray-Curtis index (Bray and Curtis, 1957; Tang et al., 2016; Silverman et al., 2016). In the context of TCR repertories, the abundance counts of observed sequences’ can be considered to be analogous to species abundance’ in an ecological context. However, because of the sparsity of observed sequences, application of these dissimilarity metrics to a dataset of TCR repertoires may not always work well. PA models have attracted substantial attention as methods to overcome these issues, since these models can compensate for the sampling bias by estimating the statistical parameters directly from the measures of the abundance counts of observed sequences (Robinson and Smyth, 2007; Sepúlveda et al., 2010), and are thus expected to be more robust to sampling bias. However, both methods ignore detailed information of the amino acid sequences of TCRs.

Therefore, in this study, rather than focusing on observation counts, we instead focus on the sequence similarity among repertoires. Our method consists of four steps: (i) calculate a dissimilarity matrix of observed TCR sequences in all samples using the Smith-Waterman (SW) algorithm with a scoring matrix; (ii) embed the data in a low-dimensional Euclidian space by dimensionality reduction methods while preserving the inter-sequence relations quantified by the dissimilarity matrix; (iii) estimate the sequence distributions in the low-dimensional space by the KDE algorithm; and (iv) quantify the sample differences by calculating the JSD value between the probabilistic density functions of different samples, and cluster the samples accordingly. Each of the above steps is described in detail in the following subsections.

#### 2.2.1 Quantification of sequence dissimilarity

The first step of our method is the quantification of similarity for each pair of TCR sequences in all samples. The SW algorithm remains the most popular pairwise local sequence alignment algorithm in bioinformatics for quantifying the similarity of amino acid sequences (Smith and Waterman, 1981). In recent years, improved versions of the SW algorithm have been proposed to resolve the problems related to the increase in computational costs along with the rapidly increasing size of datasets that are now possible from next-generation sequencing. Here, we used one of these modifications, the striped SW algorithm (Farrar, 2007), which uses a single-instruction-multiple-data (SIMD) system that allows for multiple units to simultaneously execute the same operation. The algorithm was implemented with Parasail, an open-source software for sequence alignment (Daily, 2016).

The SW algorithm requires amino acid substitution matrices, which determine the cost of the replacement of a single amino acid residue by another (Henikoff and Henikoff, 1992). Although the SW algorithm has already been applied to TCR sequences as a mapping tool for CDR3 sequences (Shugay et al., 2014), no study has yet established the best choice of substitution matrices for comparison of TCR data. Therefore, to clarify the effect of the type of substitution matrix employed and determine the optimal choice for our method, we tested 10 different matrices: five different point-accepted mutation matrices (PAM; 30, 100, 120, 160, and 250) (States et al., 1991), and five different blocks substitution matrices (BLOSUM; 45, 50, 62, 80, and 100) (Henikoff and Henikoff, 1992). The gap opening and extension penalties were set to 10 and 1, respectively (Farrar, 2007).

Since the substitution matrices give nonzero values for replacements between the same amino acid residues, the total score of the alignment between two identical sequences depends on their sequence lengths. Thus, the diagonal elements of a pairwise distance matrix will have different values even when they are calculated from the alignments of two identical sequences. In other words, both the sequence similarity and the sequence length determine the values of the pairwise distance matrix. To adjust for this sequence-length effect, we converted the pairwise distance matrix into a dissimilarity matrix using the following equation:

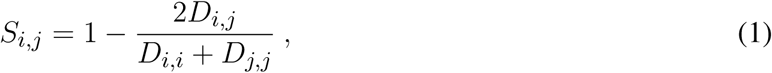

where *D*_*i,j*_ and *S*_*i,j*_ are a pairwise distance matrix and dissimilarity matrix between the two sequences *i* and *j*, respectively. At this step, we calculated the pairwise distances between all pairs of unique sequences observed in all samples with the striped SW algorithm. We then transformed the pairwise distance matrix into the dissimilarity matrix using Eq. 1.

#### 2.2.2 Dimensionality reduction with manifold learning methods

To visualize the structure of the high-dimensional dissimilarity matrix in a low-dimensional space, we applied dimensionality reduction (manifold learning) techniques to the dissimilarity matrix described above that was constructed with BLOSUM62. Here, we compared the results calculated with four different methods: multidimensional scaling (MDS) (Borg and Groenen, 2005), ISOMAP (Tenenbaum et al., 2000), spectral embedding (SE) (Belkin and Niyogi, 2001), and t-distributed stochastic neighbor embedding (t-SNE) (Van Der Maaten and Hinton, 2008). All of these methods transform the dissimilarity matrix *S* with dimensionality *N* into a new dataset *Y* with a lower dimensionality *d* in such a way as to preserve the structure of the dissimilarity matrix by minimizing cost functions. The major difference among these methods is the cost function, which is determined according to the relative distances between all pairs of sequences. MDS with a SMACOF algorithm minimizes the sum of squared errors in the relative distances of all sequence pairs before and after projections (Borg and Groenen, 2005). This cost function of MDS tends to preferentially retain the distances between more distant data points over those between more adjacent points (Van Der Maaten et al., 2009). ISOMAP also minimizes the sum of squared errors, but rather than using the relative distances, it uses the geodesic distances, which are the distances along the shortest paths between two nodes on the neighborhood graph, calculated with a k-nearest neighbor algorithm (Tenenbaum et al., 2000). In the present study, we calculated the geodesic distances with the Warshall-Floyd algorithm (Floyd, 1962). ISOMAP retains a neighborhood structure of data points lying on a curved manifold (e.g., the Swiss roll dataset (Tenenbaum et al., 2000)), which is collapsed in MDS. SE, also known as Laplacian eigenmaps, minimizes the cost function based on the neighborhood graph, which ensures that local neighborhood relations in a high-dimensional space are preserved in a embedded low-dimensional space (Belkin and Niyogi, 2001, 2003). We regarded the adjacency matrix based on the k-nearest neighbor algorithm as the weighted graph matrix to construct the Laplacian graph of SE. Finally, t-SNE converts the relative distances to joint probabilities, and minimizes the Kullback-Leibler divergence between the joint probabilities of the high-dimensional space and those of a embedded low-dimensional space (Van Der Maaten and Hinton, 2008). For calculation of the joint probabilities, t-SNE uses different kernels for the high-and low-dimensional spaces: a Gaussian kernel and a Student t-distribution, respectively. Since the Student t-distribution results in heavier tails than the Gaussian kernel, the t-SNE method emphasizes the local distances between data points in the low-dimensional space.

In the studies of sequence alignments for sequences with different lengths, it is impossible to know the precise coordinates and the dimension of the sequence space. Thus, we cannot directly use principal components analysis, which is the most widely used dimensionality reduction technique (Bishop, 2007) but requires vector data with fixed dimensionality. The common advantage of the above four methods is that if the distances between all pairs of data points are known, then there is no need to know the specific coordinates of the sequence space (Van Der Maaten and Hinton, 2008; Van Der Maaten et al., 2009).

We implemented t-SNE, MDS, and SE with the Scikit-learn manifold learning library with Python (Pedregosa et al., 2012). ISOMAP was implemented with our custom-written code in Python, because the ISOMAP function of the Scikit-learn toolbox does not support the dissimilarity matrix as an argument. The detailed parameters of all methods are described in Table S1 in the Supplementary Information.

#### 2.2.3 Estimation of the probability density function with KDE

To compare the data points scattered in the embedded low-dimensional space among different samples, the embedded discrete data can be interpolated with a probability density function (PDF). Here, we estimated the PDF with the KDE algorithm (Jones et al., 1996; Heidenreich et al., 2013; Arlot and Celisse, 2010). The exponential function was used as the kernel of the KDE (Christopher et al., 1997). The bandwidth parameter of the exponential kernel function was optimized by maximum-likelihood estimation with a cross-validation algorithm (Arlot and Celisse, 2010). To reduce the computational cost of this calculation, we utilized the Kd-tree algorithm, which is an N-body algorithm that divides all of the data into N clusters based on their relative Euclidean distances (Gray and Moore, 2001). KDE was implemented with the parameter optimization toolbox in Scikit-learn (Pedregosa et al., 2012). For application of the KDE, we discretized the embedded space with 400 bins along each axis with the following range: [min *x*_*i*_- (max *x*_*i*_- min *x*_*i*_)/10, max *x*_*i*_ + (max *x*_*i*_- min *x*_*i*_)/10], where *x*_*i*_ indicates the position of a data point (i.e., a sequence) in the embedded space, and *i* indicates each axis of that space.

#### 2.2.4 Quantification of sample differences with JSD

The final step of our method involves quantification of the inter-sample differences by calculating the JSD values between all pairs of the estimated PDFs (Elhanati et al., 2014). The JSD is defined as:

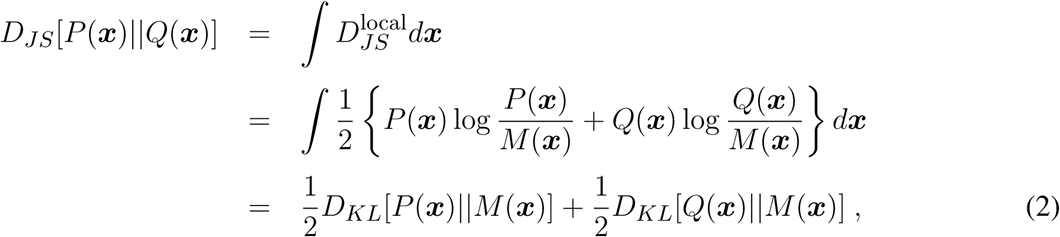

where *P* (***x***) and *Q*(***x***) are the estimated PDFs and *D*_*KL*_ is the Kullback-Leibler divergence; *M* (***x***) is 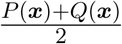 and 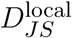 is the “local JSD”, whose integration with respect to ***x*** gives the JSD. Thus, the “pairwise” JSDs provide a sample-difference matrix that quantifies the combinatorial differences between all pairs of the samples. To categorize all samples, we utilized hierarchical cluster analysis, which converts the *N* × *N*-dimension sample-distance matrix into a dendrogram. Specifically, we used an agglomerative hierarchical cluster technique; each sample is initially treated as a singleton cluster, and pairs of clusters are repetitively merged according to a criterion until only a single cluster remains (Maimon and Rokach, 2005).

We here used Wards criterion (Ward, 1963) for agglomerative clustering, which was implemented using the linkage function of Matlab’s Statistics and Machine Learning Toolbox (The MathWorks Inc., Natick, MA, USA). To compare our clustering result of the observed sequences with those obtained using other count-based methods, we also quantified the inter-sample difference with BPLN and Bray-Curtis methods. BPLN was applied according to the methods described in the original paper by Rempala et al. (2011).

To evaluate the goodness of fit of the clustering results, Rempala and colleagues Rempala et al. (2011) calculated the cophenetic correlation coefficient (CCC), which quantifies the distortion due to the transformation from the distance matrix to the cophenetic matrix, from which the dendrogram was derived. However, the CCC does not always accurately reflect the goodness of fit of the results. Indeed, Wards method tends to produce lower CCC values than other methods such as average and centering methods even though it has been previously reported as the best agglomerative method (Hands and Everitt, 1987; Saracli et al., 2013). Therefore, instead of the CCC, we verified the fit of the model based on the statistical significance of the distance between the nodes of the dendrogram, because the significance of the estimated value of JSD is unclear. Specifically, we used bootstrap methods to evaluate significance, resampled data points from the naive PDF according to the number of observed read counts, and then re-estimated the PDF from the resampled data points. We then calculated the JSDs between the naive and re-estimated PDFs. We repeated this process 100 times to obtain a histogram of the calculated JSDs. The 2.5th and 97.5th percentiles of the histogram of the JSDs between the naive and each re-estimated PDF represent both ends of the 95% confidence interval, where values outside of the interval indicate a significance level of over 5%.

To identify the sequences with the greatest contributions to the inter-sample distances, we selected square bins for the top 1% of the local JSDs. We next defined the sequences in these bins as those contributing to the observed pairwise sample difference. Furthermore, to investigate the characteristics of the contributing sequences, we calculated the relative frequencies of the amino acid residues in all of the contributing sequences. The graphics of the relative frequencies were obtained using WebLog 3 software (Crooks et al., 2004).

All analyses were performed using custom-made codes written in Python, Matlab, and R.

## 3 RESULTS

### 3.1 Evaluation of sequence dissimilarity for pairwise sequence alignment

Using the pairwise sequence alignment and Eq. 1, which excludes the influence of the sequence lengths from the alignment results, we calculated the dissimilarity matrix of all pairwise sequences in the dataset of Rempala et al. (2011). The upper panels in Figs. 1(A) and 2(B) show the dissimilarity matrices obtained with the 10 different substitution matrices, five of PAM and five of BLOSUM. As shown in these panels, the components of the dissimilarity matrices are clearly separated into two distinct clusters, which are considered to reflect the *α*-chain rearrangements between J*α*2.6 and J*α*2 under the usage of the other fixed VJ genes. Moreover, the low-numbered PAMs and high-numbered BLOSUMs showed more gradual differences among the matrix elements than the others. This tendency was even more evident when viewing their embedded spaces for the separation of clusters. The lower panels in Fig. 1(A) and (B) show the t-SNE projection maps of the corresponding dissimilarity matrices in the upper panels. In this case, low-numbered PAMs and high-numbered BLOSUMs tended to have merged clusters. This may be attributed to the specific characteristics of these two substitution matrices, which have a higher variability in the scores for replacements between a pair of amino acids. Based on these results, we used the BLOSUM62 dissimilarity matrix for subsequent analyses for two main reasons. First, both the too high-numbered PAMs and too low-numbered BLOSUMs seemed to lose the intra-cluster structures by trying to compress the clusters into regions that were too small, while both the too low-numbered PAMs and too high-numbered BLOSUMs diminished any inter-cluster differences, resulting in indistinguishable clusters. Second, BLOSUM62 has been the most widely used matrix in analyses of TCRs and antigen peptides to date (Oyarzún et al., 2013; Schwaiger et al., 2014; Hoffmann et al., 2015; Aas-Hanssen et al., 2015).

**Figure 1.**
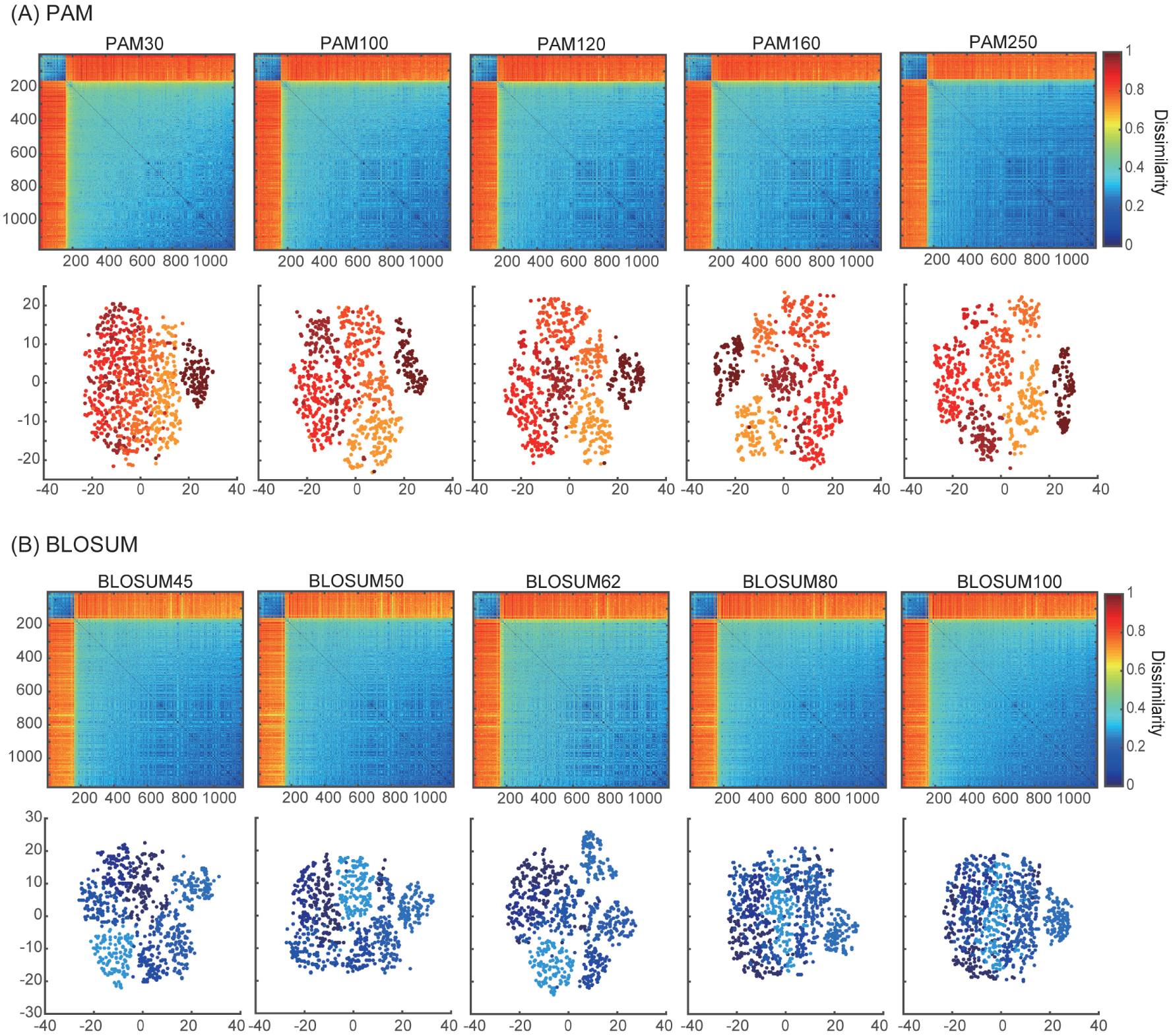
Dissimilarity matrices and their embedded distributions with 10 different score matrices; (A) PAM, (B) BLOSUM. The upper and lower panels show the dissimilarity matrices and projection maps in two-dimensional space, respectively. All of the rows and columns in each dissimilarity matrix were sorted according to the sum of their elements. The colors of points in the lower panels of (A) and (B) correspond to the clusters in PAM250 and BLOSUM45 that were discriminated by k-means algorithms (k = 7).

**Figure 2.**
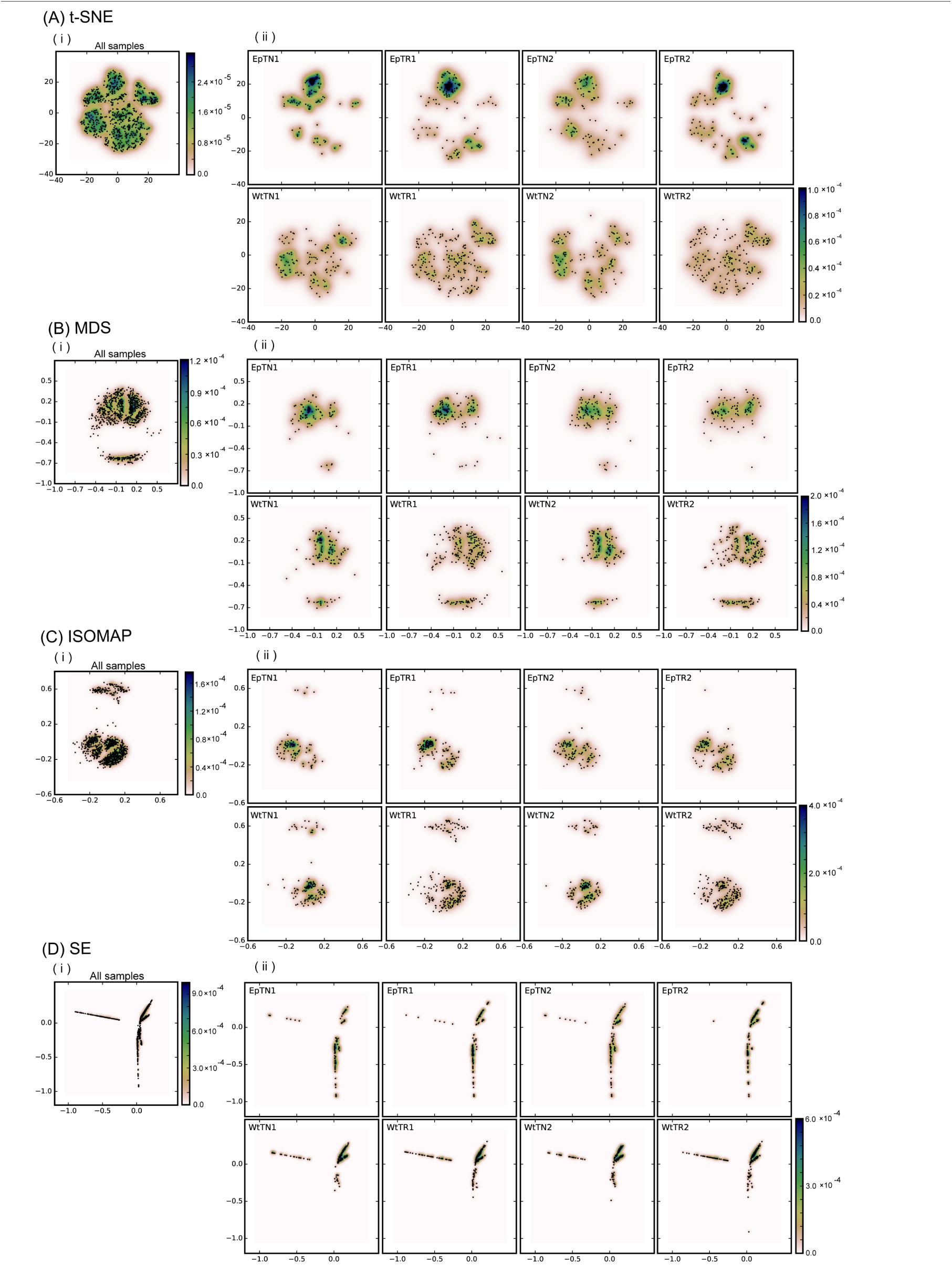
Dimensional reduction with four different dimensionality reduction methods: (A) t-SNE, (B) MDS, (C) ISOMAP, and (D) SE. Panel (i) includes the points of the total unique sequences observed in all samples. Panel (ii) includes only the portions of sequences that were observed in each sample. “Ep” and “Wt” denote two different genetic backgrounds of mice. “TN” and “TR” denote naive and regulatory T-cells. “1” and “2” denote the thymus and peripheral lymph nodes, respectively. As an instance, EpTN1 denotes the naive T-cells that were collected from the thymus in the “Ep” mice.

### 3.2 Dimensionality reduction of the dissimilarity matrix

To evaluate the applicability of dimensionality reduction methods, we reduced the dimensionality of the dissimilarity matrix into a two-dimensional space using four different dimensionality reduction methods (t-SNE, MDS, ISOMAP, and SE). In Fig. 2, each point in each panel corresponds to a unique sequence of TCRs, and the spatial distances between pairs of points reflect the dissimilarity of the sequences corresponding to the points. Panels (i) and (ii) of Fig. 2(A–D) show the projection results of the unique sequences obtained from all samples, and the subset of points (sequences) in (i) that appeared in the sample denoted in the inset letters of the panel, respectively. The sample differences could be clearly reflected according to the scattering pattern of the points. Moreover, the points derived from the t-SNE and MDS methods spread more widely over the two-dimensional space than the others, whereas the points were more locally consolidated with the ISOMAP method, and especially with SE. This result suggests that t-SNE and MDS may be more appropriate than other reduction methods for larger datasets, because highly dense regions can cause difficulty in comparing the probabilistic distributions between samples. Furthermore, the two clear clusters in the dissimilarity matrix (the upper panel of BLOSUM62 in Fig. 1(B)) were well reflected in the two clusters for the MDS and ISOMAP methods (Fig. 2(B,C)), but were not represented clearly in the clusters of t-SNE. This result suggests that t-SNE emphasizes slight differences within clusters rather than large differences between the clusters of the dissimilarity matrix. Since it is unclear whether this visualization property of t-SNE works efficiently for comparisons between samples, we quantified and compared the distributions of data points at the next step, and examined the method that would be most appropriate for this purpose.

### 3.3 Hierarchical clustering of the pairwise-sample-difference matrix

We applied the KDE algorithms to the spatial distributions of data points to estimate their probability density functions (color gradient in Fig. 2), with which JSD is calculated to quantify pairwise-sample differences. The matrices of the pairwise-sample differences are shown in Fig. 3(A), and the resulting dendrograms in Fig. 3(B) indicate the hierarchical clustering results with the agglomerative method. The clustering results can be categorized into two groups: ISOMAP and the others (MDS, t-SNE, and SE). The dendrograms of t-SNE, MDS, and SE showed good correspondence with our intuitive notions about the hierarchical structure of the experimental conditions, which were ranked in order of donor sites, types of T cells, and genetic background with clear biological significance (Rempala et al., 2011). By contrast, the dendrogram of ISOMAP showed a mismatch in the hierarchical order between the T-cell types and donor sites of Ep mice.

**Figure 3.**
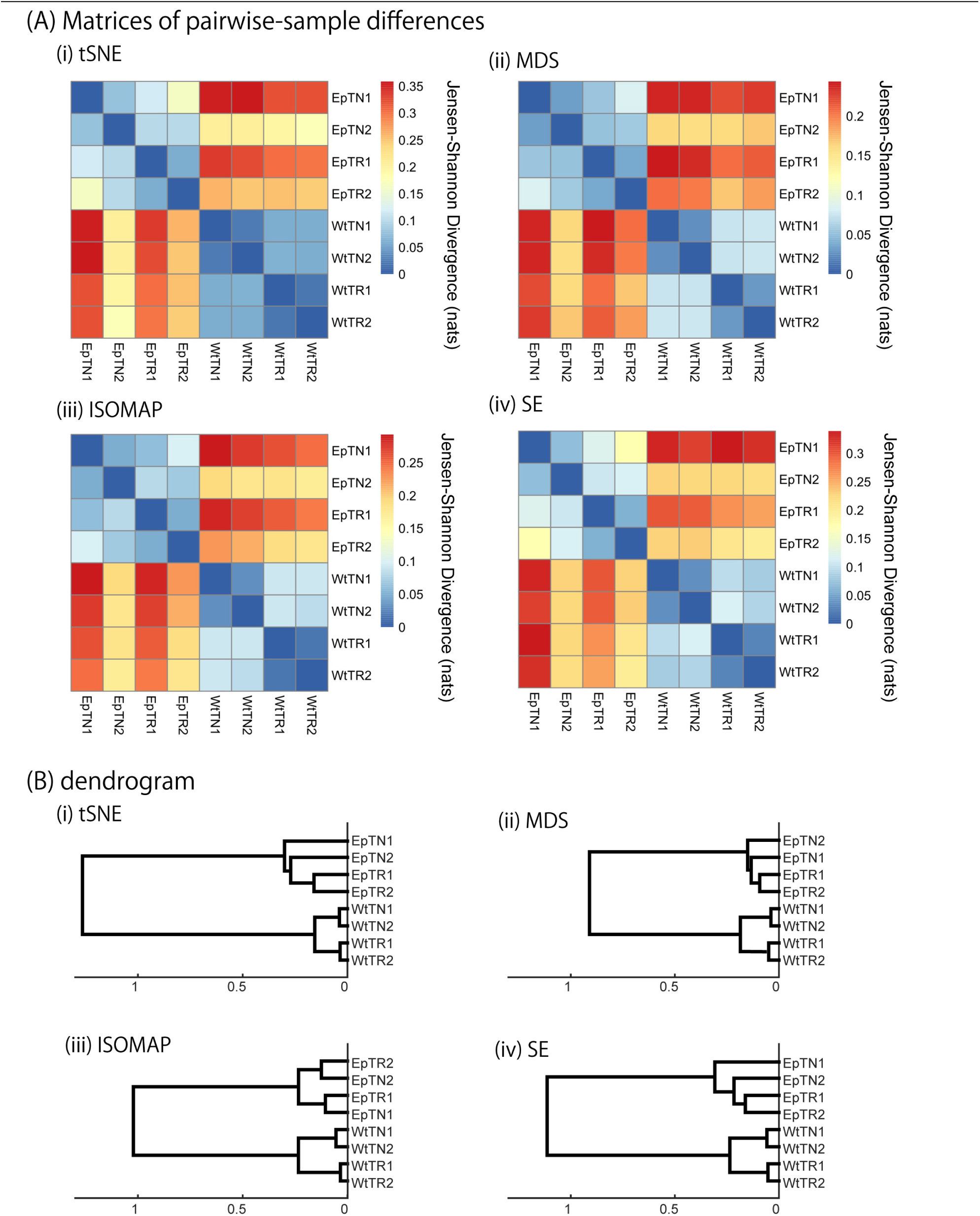
JSD matrices and their clustering results with four different methods: (i) tSNE, (ii) MSD, (iii) ISOMAP, and (iv) SE. (A) Matrices of pairwise-sample differences and (B) the dendrogram constructed from the matrices.

To verify the results of hierarchical clustering obtained by our method, they were compared with those obtained with previous observation count-based methods, the BPLN and Bray-Curtis method. As shown in Fig. 4, the sample differences and dendrograms estimated from the BPLN and Bray-Curtis methods were very similar to those obtained using our approach with MDS, t-SNE, and SE. Importantly, these similar results were obtained with different data modalities: sequence similarity and observation counts. Therefore, this consistency suggests that there is common information between sequence similarity and observation counts with respect to quantifying the differences among samples. We should note that these two modalities can be combined simply by assigning the number of observed sequence counts as a weighting factor for each data point (i.e., a unique sequence) in the embedded space. Indeed, the counts-weighted PDFs using KDE (Fig. S1) showed no obvious change in the hierarchical clustering structure of the pairwise-sample differences. Taking these results together, the MDS or t-SNE appears to be the better choice as a dimension-reduction method for evaluation of differences in TCR repertories among samples, given that these methods show wide spatial distributions of the data points, and also show the most consistent dendrogram structures with those of previous count-based methods.

**Figure 4.**
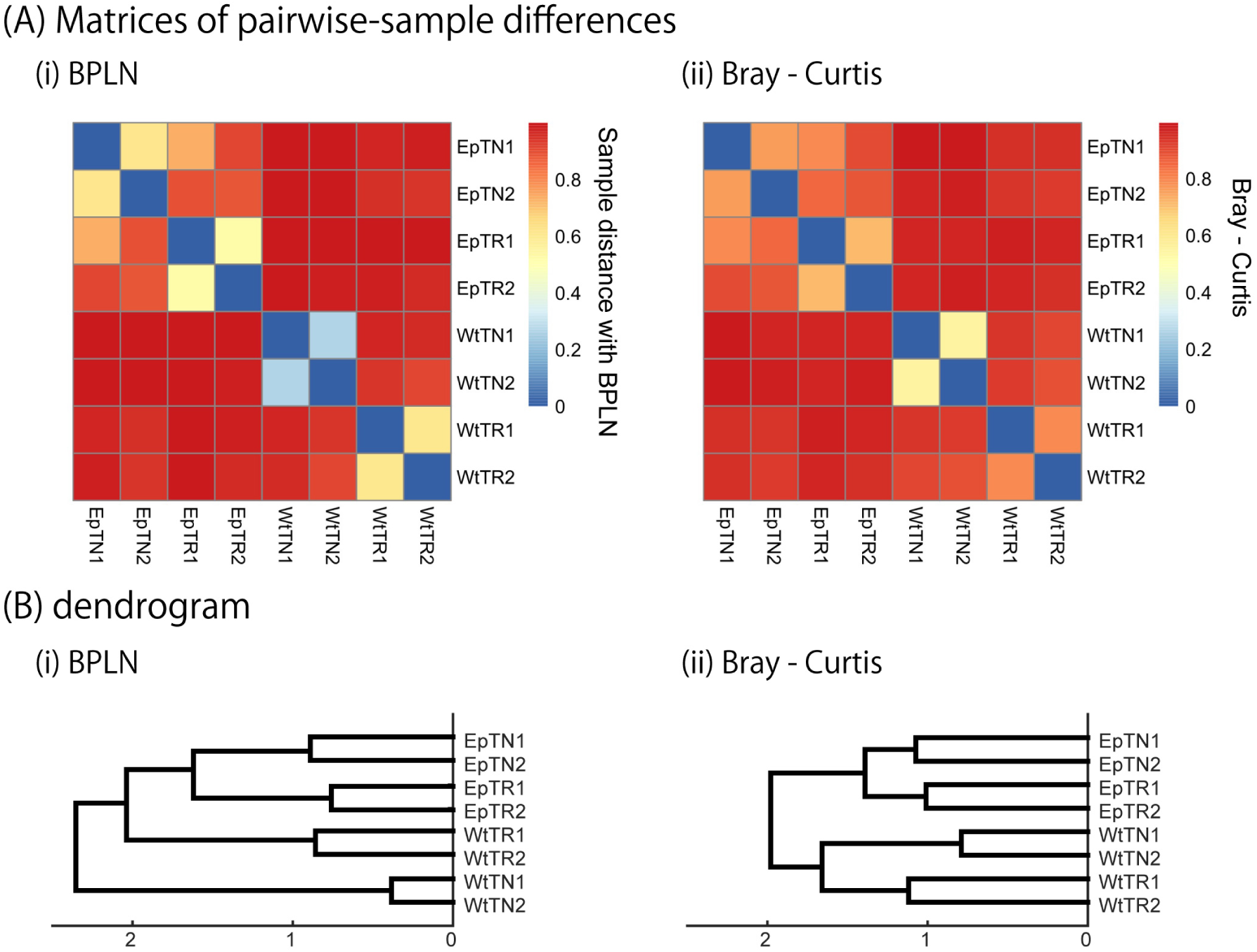
Sample-distance matrices constructed with two methods: (i) BPLN, (ii) Bray-Curtis. (A) Matrices of pairwise-sample differences and (B) the dendrogram constructed from the matrices.

### 3.4 Significance test for inter-sample differences

To verify the statistical significance of the calculated JSDs between all sample pairs, we calculated the JSDs between the naive and the re-estimated PDFs using a non-parametric bootstrap algorithm. Figure 5 shows the histogram of the JSD values between the naive PDF of Fig. 2 (C, ii) and the re-estimated PDFs. In the figure, arrows indicate the naive JSD values between the sample designated on the top of the panel and the other samples. If the values indicated by the arrows are bigger than the light red region in the panel, the pairwise naive PDFs deriving the naive JSD are significantly different each other. Except for EpTN2, the arrows that indicate the JSD values between the pairs in proximity to the terminal nodes of the dendrogram in Fig. 3 (A) were in inside of the light red regions, which means that the JSD values were not significantly bigger than the JSD values of the histogram. This result suggests that the PDFs of these pairs are so similar that they cannot be statistically distinguished from each other. By contrast, the arrows that indicate the naive JSD values between pairs in the upper parts of the clusters, above the terminal nodes, were in outside of the red light regions, which means that the JSD values were significantly different from each other. This result indicates that the types of T cells and the genetic background can be discriminated with sufficient statistical significance.

**Figure 5.**
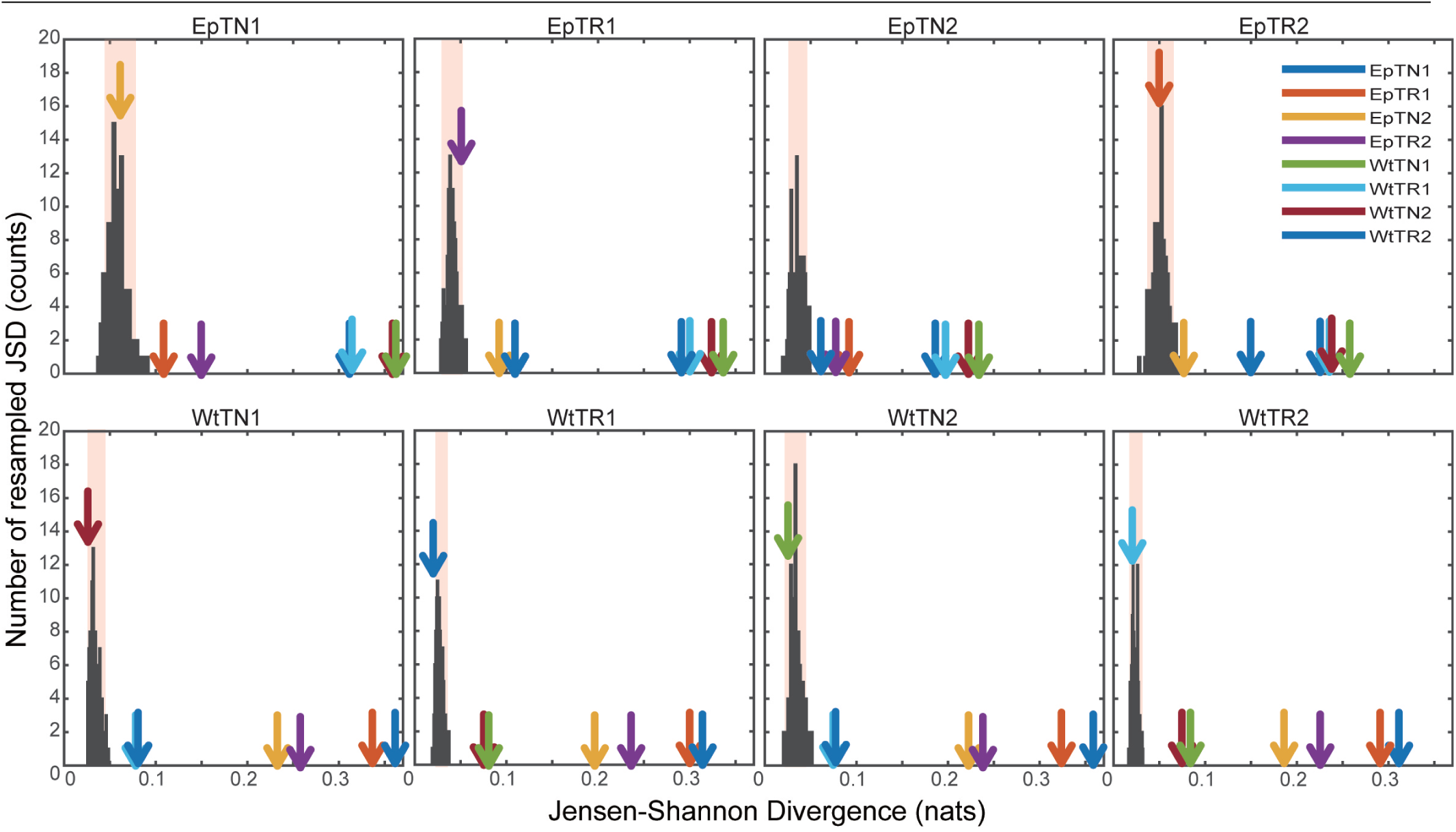
Significance tests of JSD values using bootstraps. Each colored arrow indicates the naive JSD values between the resampled sample and another. The light red region indicates the 95% confidence interval.

### 3.5 Spatial distribution of local JSD values

The main advantage of our method compared to count-based methods is the ability to identify the major sequences contributing to inter-sample differences. To identify the sequences with the greatest contributions to large local JSD values, we plotted the spatial distribution of the local JSDs between the WtTN1 and EpTN1 sequences. As shown in Fig. 6, two regions were identified that were associated with top 1% significance values. Table 1 lists the identified sequences in these regions with larger local JSDs than the others. In regions 1 and 2, there were no sequences for WtTN1, whereas EpTN1 had multiple sequences in these regions, suggesting that these apparent Ep-specific sequences may contribute to the observed abnormality in the antigen presentation of Ep mice.

**Figure 6.**
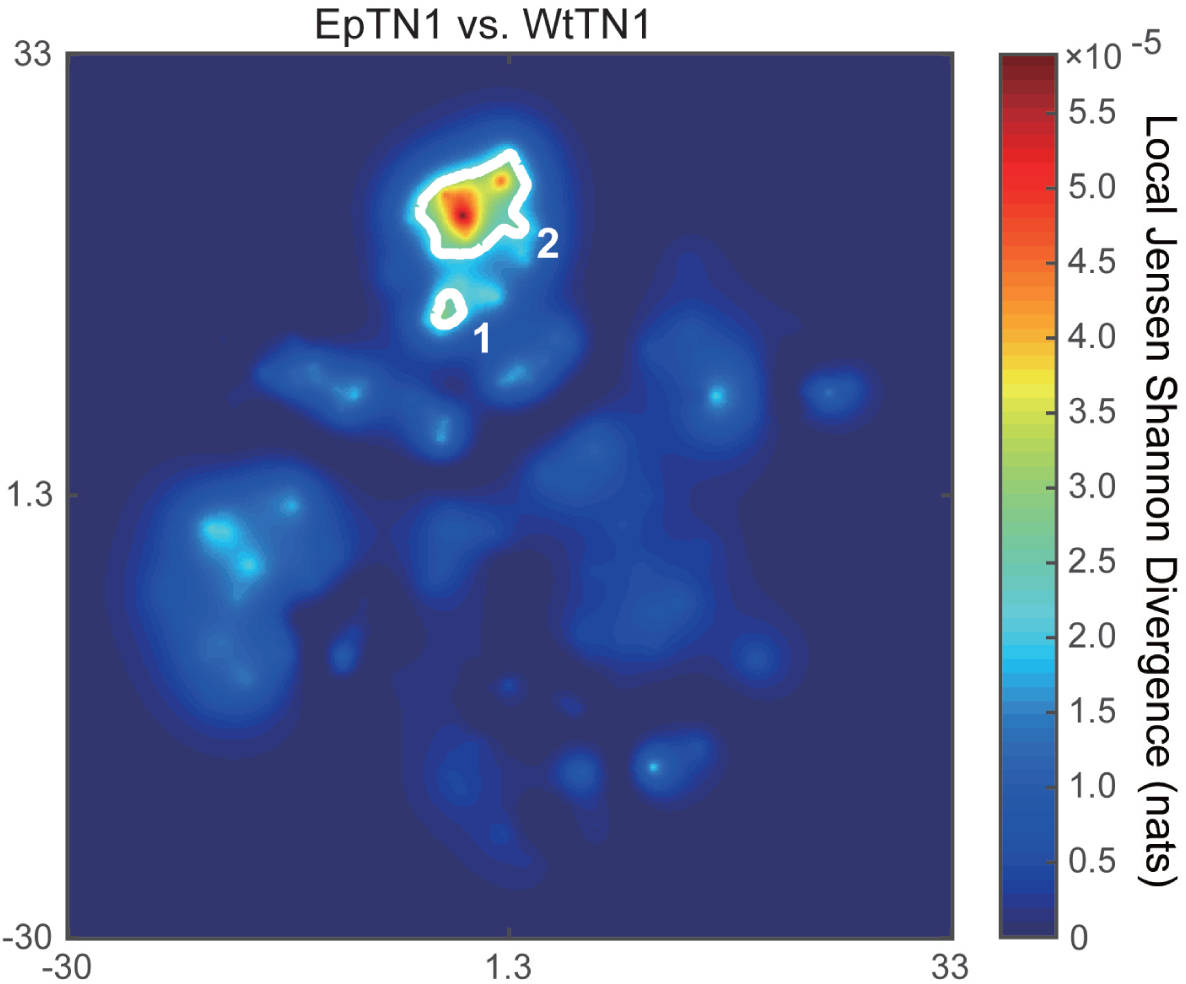
Spatial distribution of local JSD values between EpTN1 and WtTN1. The white line shows the contours of the regions with significant local JSDs.

**Table 1.** Sequences contributing to the JSD between EpTN1 and WtTN1.

| Regions | EpTN1 | WtTN1 |
| --- | --- | --- |
| 1 | CAASAYQLIWG<br>CAASCYQLIWG<br>CAASRYQLIWG<br>CAASTYQLIWG |  |
| 2 | CAAGNYQLIWG<br>CAAHNYQLIWG<br>CAANNYQLIWG<br>CAARNYQLIWG<br>CAASNYQLIWG<br>CAATNYQLIWG<br>CADLNYQLIWG<br>CADSNYQLIWG<br>CAGSNYQLIWG<br>CASHNYQLIWG<br>CASSNYQLIWG<br>CATSNYQLIWG<br>CAVSNYQLIWG<br>CVGSNYQLIWG |  |

This type of sequence identification can provide further knowledge about the characteristics of sequence alignments. Figure 7 shows the occupation probability (relative frequency) of amino acids at each position of the sequences obtained from all of the sequences contributing to pairwise differences, and a consensus sequence was determined from amino acid positions 6th to 11th. We note that these contributing sequences and their characteristics cannot be easily identified simply by examining the overlapping sequences in two samples, because there was almost no overlap between EpTN1 and WtTN1 sequences (0.352%, 1/284), and because these account for only 6.34% (18/284) of the total unique sequences in the two samples.

**Figure 7.**
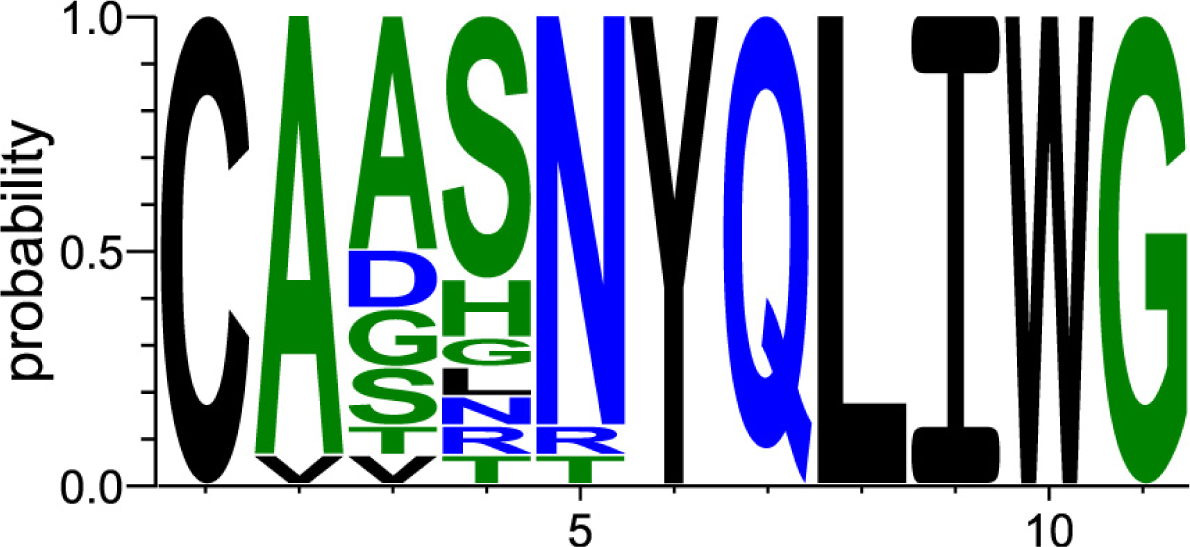
Relative frequencies of observed amino acids at each position in the contributing sequences.

## 4 DISCUSSION

We quantified the difference in TCR repertoires among different samples based on amino acid sequence dissimilarity. Through a quantitative comparison of the sequence distributions in the dimension-reduction spaces of the dissimilarity matrix, we estimated the inter-sample hierarchical structure, which was almost identical to that estimated with previous count-based methods that did not incorporate detailed sequence information. Furthermore, we identified the sequences that most strongly contribute to the pairwise sample difference using the local JSD distribution.

Despite the fact that our method relies on sequence similarity and previous methods are based on observation frequency, completely different features of the TCR repertoire, almost the same sample-clustering structure was obtained. This suggests that there is a relationship between the observation counts and sequence similarities, which was further confirmed by the lack of change in the structure of the hierarchical clustering result when estimating the PDFs by the KDE algorithm with the weights based on the number of observed sequence counts. Further studies to understand this relation in greater depth would allow to cross-check the results of sample classification by investigating the consistency of two methods. Moreover, clearly distinguishing the overlapping and non-overlapping information between counts and sequences may allow for more detailed classification of samples.

Although our method and the counts-based methods provide similar classification result, there are two unique merits of our method. The first is the robustness against errors derived from polymerase chain reaction (PCR) amplification bias attributed to the variability in reproducibility for individual sequences (Greiff et al., 2015). Previous studies have shown that the PCR efficiency is affected by sequence profiles such as the length and GC content (Aird et al., 2011; Kivioja et al., 2011). Indeed, high-throughput sequencing with DNA barcoding has confirmed that the PCR amplification efficiency of TCR sequences is highly variable due to the differences in profiles of individual cDNA molecules (Carlson et al., 2013; Shugay et al., 2014; Best et al., 2015). This fact suggests that the PCR process for TCR sequence amplification induces errors in the numbers of observed sequence counts, which may eventually lead to errors in the results of counts-based methods such as PA models. Alternatively, our method does not depend on the sequence counts, allowing for reliability against errors due to PCR bias. The second key merit of our method is the ability to identify the sequences with the greatest contributions to pairwise sample differences. This sequence identification allows for targeted analyses along with the results of other studies such as the simulation modeling for determining the crystal structures of the TCRs encoded by these sequences (Klausen et al., 2015) or establishing alignments between CDR3 sequences and microbial genomes (Aas-Hanssen et al., 2015). Such a closed-loop experimental design may help to achieve a breakthrough in the development of vaccines or immunotherapies (Hou et al., 2016). Thus, the present results and advantages demonstrate the potential applicability of adopting a sequence-based method in repertoire analysis, which can compensate for the drawbacks in conventional count-based methods.

Nevertheless, there are several issues and problems that should be mentioned that are worthy of further investigation in developing and improvement of sequence-based approaches for comparison of TCR repertoires.

One issue concerns the treatment of gap penalties. When we evaluated the differences of the score matrices shown in Fig. 1, we fixed the gap opening and extension penalties to 10 and 1, respectively. Although the effects of the penalties have not been adequately investigated in previous studies (Wrabl and Grishin, 2004), the gap opening penalty was found to affect estimations of the hierarchical data structure (data not shown). The CDR3 region of TCR is a much shorter sequence than peptide sequences, and also shows frequent deletions and insertions from somatic recombination events. Considering these characteristics, further investigations about the effects of gap penalties are needed.

Another aspect worthy of further consideration is the empirical estimation of the cost functions used in the dimensionality reduction methods and in the combination of projection methods and comparison of the embedded results. As demonstrated in Fig. 2, the scattering patterns of the sequence data in the low-dimensional space depend on the cost functions of the method adopted. Using MDS and ISOMAP, we obtained two clear clusters reflecting two regions in the dissimilarity matrix. This is because the cost function of MDS preferentially preserves the distances between the distant points rather than those between nearby points (Van Der Maaten et al., 2009). By contrast, t-SNE emphasizes the local structures of nearby points over global points by using the Student t-distribution as the kernel of the embedded space. These cost functions were empirically determined for visualization purposes in the original papers, without consideration of the subsequent quantitative inter-sample comparison of the embedded results. Although our results suggest that the empirical combination of dimensionality reduction methods and comparison of the embedded results by JSDs may work well, both the projection method and comparison method in the embedded space should be consistently designed so as to best reflect the inter-sample difference in the original sequence space. This method might be developed by choosing an information-theoretic measure for the cost function of projection that can preserve the relevant information of repertoires in the original sequence space. Because the underlying high-dimensional structures of the repertoire are difficult to capture intuitively, methods based on firm theoretical rationality and biological significance are indispensable for further exploitation of repertoire information.

## DISCLOSURE/CONFLICT-OF-INTEREST STATEMENT

The authors declare that the research was conducted in the absence of any commercial or financial relationships that could be construed as a potential conflict of interest.

## AUTHOR CONTRIBUTIONS

R.Y., Y.K., and T.J.K: study conception and design; R.Y. and Y.K.: performed the research; R.Y. and Y.K: data analysis; and R.Y. and T.J.K.: wrote the paper.

## FUNDING

This research is partially supported by the Platform Project for Supporting in Drug Discovery and Life Science Research (Platform for Dynamic Approaches to Living System) from the Japan Agency for Medical Research and Development (AMED), Grant-in-Aid for Exploratory Research (15K14433) from the Ministry of Education, Culture, Sports, Science, and Technology, Japan, and JST PRESTO Grant Number JPMJPR15E4, Japan.

## ACKNOWLEDGMENTS

We are grateful to Shohei Hori, Saito Yohei, Yotaro Katayama, and Kazumasa Kaneko for fruitful discussions.

## SUPPLEMENTARY INFORMATION

**Figure S1.**
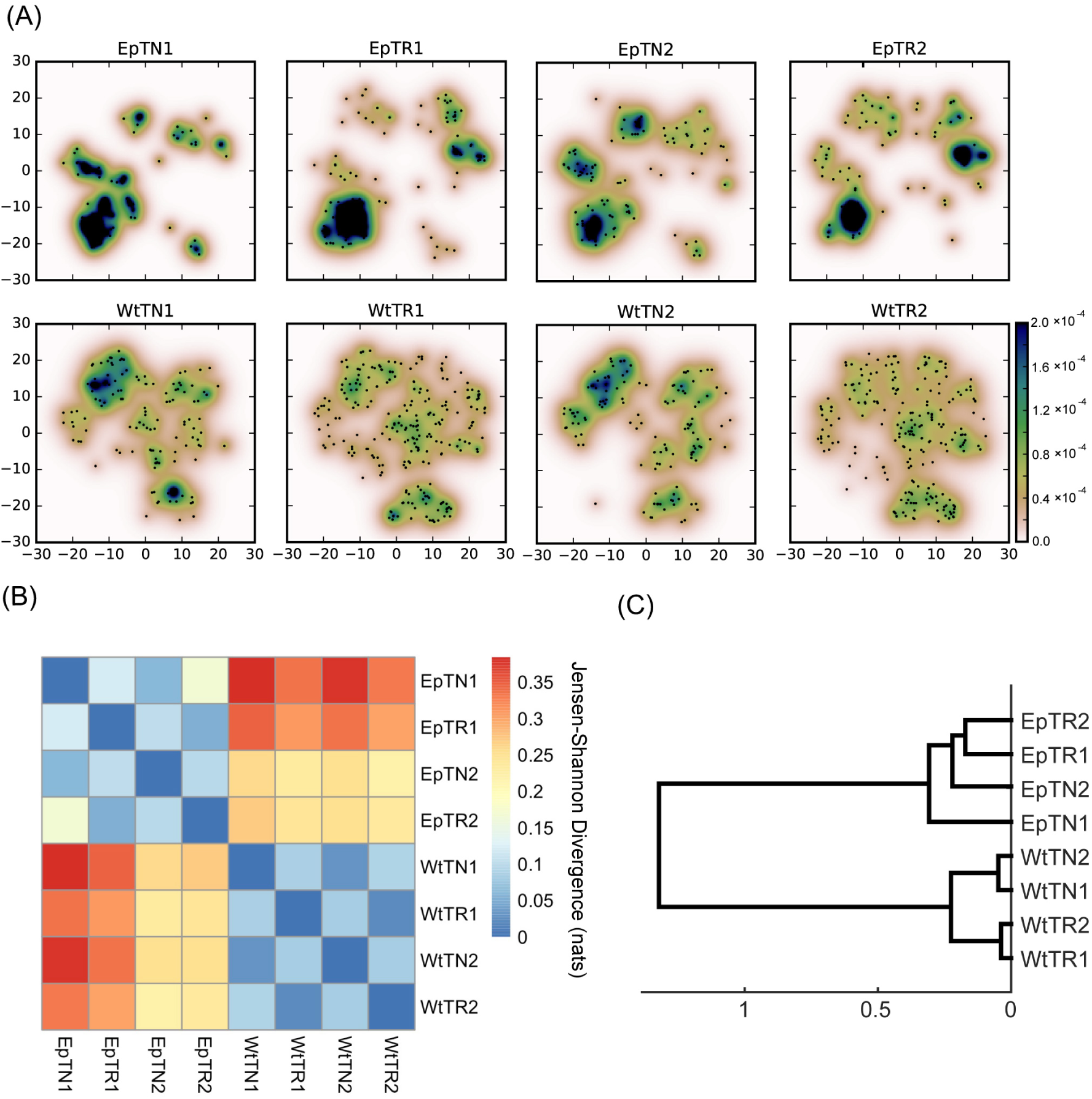
Count-weighted PDF estimation with tSNE-embeding data. Conventions comply with Fig.2 and 3. (A) PDFs weighted by the number of sequence counts. (B) Sample distance matrix estimated with the weighted PDFs. (C) the dendrogram constructed from the matrix in (B).

**Table S1.**
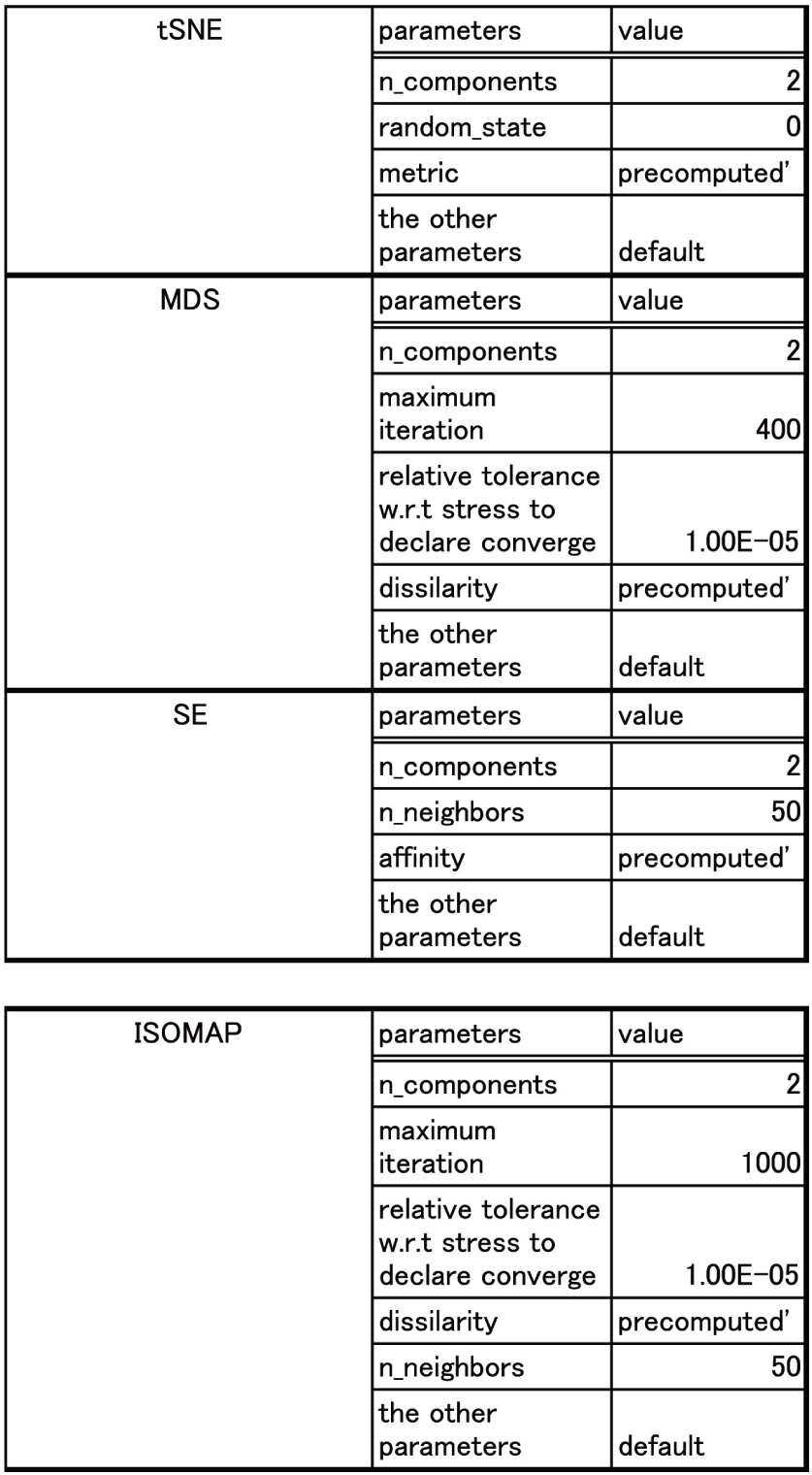
Detail parameters of manifold learning methods. Except for ISOMAP, we used the functions in the class of sklearn.manifold in the scikit-learn toolbox. The parameters of each function are described in the Table S1. For ISOMAP, first, we calculated the geodesic distances by using k-nearest neighbor algorithm and Floyd-Warshall method. Then, we applied the geodesic distances to MDS with the following parameters.

